# Modulation of Pain Perceptions Following Treadmill Running with Different Intensities and The Potential Mechanisms of Exercise-induced Hypoalgesia

**DOI:** 10.1101/2023.04.17.537131

**Authors:** Zi-Han Xu, Nan An, Jeremy Rui Chang, Yong-Long Yang

## Abstract

**Objective:** This study aimed to compare the effects of three intensities of treadmill running on pain perceptions in healthy individuals. And investigate the role of endogenous pain modulation in the exercise-induced hypoalgesia (EIH) effects.

**Methods:** Sixty-six healthy female individuals were included in this study and were randomly assigned to one of three treadmill running intensities for 35 minutes: 40% of their reserve heart rate (HRR), 55% HRR, or 70% HRR. The EIH effects were assessed by the changes of pressure pain thresholds (PPT) and pressure pain tolerance thresholds (PPTol) at multiple time points. The assessments were conducted prior to the treadmill running session every 5 minutes during the exercise bout, and at 5 minutes, 10 minutes, and 24 hours post-exercise. The conditioned pain modulation (CPM) was also measured to determine the functions of endogenous pain modulation.

**Results:** Compared with baseline, there was a significant increase of PPT and PPTol at arm and leg in all groups during running and 5-10min follow-ups. The PPT and PPTol changes of moderate and low intensity groups were significantly higher than the high intensity group during running and 24h after running. While the CPM responses of high intensity group were significantly reduced compared with other groups at 24h follow-up.

**Conclusion:** Moderate and low intensity running may trigger the endogenous descending inhibition and elicit significant EIH effects following running and persisting over 24h. While the high intensity running only induced limited EIH effects for the activation of both descending pain inhibition and facilitation, with reduced CPM responses. Thus, the pain perception changes following exercises may reveal the potential mechanisms of EIH induced via exercises with different intensities.

**New findings:** What is the central question of this study?

Both the primary analgesia effect (EIH) and secondary pain allodynia (delayed onset muscle soreness) may occur following exercises, possibly due to the interaction between endogenous pain modulation and exercise intensities. What is the difference in the changes of primary and secondary pain perceptions following exercise with different intensities?

What is the main finding and its importance?

Moderate and low intensity running induced acute and long-lasting EIH effects via the effective activation of descending inhibition, while the high intensity running may trigger the descending facilitation and attenuate both the acute and long-lasting EIH effects. This result preliminarily explained the non-liner effect of exercise intensity on the acute EIH responds.

## Introduction

Exercise has been identified as an effective intervention to manage individuals with pain syndromes. Previous studies have demonstrated that global aerobic exercises^1^ or local resistance exercises^2^ can effectively alleviate pain perception^3^ and enhance the emotional well-being, which is commonly referred to as exercise-induced hypoalgesia (EIH)^4^.The endogenous pain modulation^5^ and the cortex cognitive process of pain^6^ are associated with the EIH, where exercise can trigger the endogenous descending inhibition to the nociceptive signals from the spinal cord dorsal horn^7^, and activate of the reward circuits of the corticolimbic systems^8^.

The analgesic effect following exercises in asymptomatic individuals tends to be correlated with exercise intensities^9,10^, while the pain tolerance thresholds can also be significantly improved during various physical activity^11^. However, the individual’s EIH can be affected by the function of endogenous pain modulation^12^ or psychological factors^13^ such as pain catastrophizing, fear, anxiety, or depression. Notably, high-intensity exercises (above 6 METs) often failed to restore the pain detection thresholds in individuals with pain conditions^14^, while low or moderate-intensity activities (3-6 METs) can still increase the pain tolerance thresholds.^15^

Differential effects of EIH effects between exercise intensities might be attributed to the mechanisms of endogenous pain modulation, which have different activation thresholds towards perception stimulus^16^. The ventromedial nucleus (VM) of thalamus can be activated by inputs from both noxious and non-noxious C fibers, which can be triggered via muscle contraction^17^, and induce the descending inhibition. And the mediodorsal nucleus (MD) of thalamus can trigger descending facilitation with inputs from noxious C fibers. Then, the neurons in periaqueductal gray (PAG)^18^ and rostral ventromedial medulla (RVM)^19^ can modulate the nociception signals through the projection to the dorsal horn of spinal cord and change the pressure, heat, or cold pain detection and tolerance threshold (PPT, PPTol, HPT, HPTol, CPT, and CPTol).

Considering the threshold of descending facilitation is relatively lower than the inhibition^20^, the activation of pain inhibition usually requires high intensity painful stimulus (conditioned stimulus), which is called conditioned pain modulation (CPM). Thus, exercise with high intensity may activate the opioid^21^ systems in the PAG and RVM^22^, then induce the pain inhibition, but may not in the individuals with impaired CPM function^23^. And the high intensity exercise may also trigger the descending facilitation^24^, which may decrease both EIH and CPM responses, lead to the mechanical allodynia^25^ and even produce the delayed onset muscle soreness (DOMS)^26^ in asymptomatic individuals^27^.

However, the moderate or low intensity stimulus may still activate the cannabinoid^28^, and 5-HT^5^ systems, and trigger the descending pain inhibition via the temporal summation^29^ of non-noxious C fibers inputs, without triggering the nociceptors and descending facilitation. And the inhibition-only EIH effects following moderate or low intensity exercise might not induce the mechanical allodynia or DOMS, which have not been verified in human studies yet.

Considering the potential modulation of EIH effect in exercises with different intensities, this study aimed to compare the changes of PPT and PPTol in asymptomatic individuals following running exercises with various intensities. And we were also investigated changes of the CPM responses before and 24h after the running sessions to reveal the potential mechanisms of EIH induced by exercises with different intensities

We hypothesized that: (1) Low-, moderate- and high-intensity running exercise might elicit EIH responses and increase the PPT and PPTol. (2) The analgesic effects of moderate or low intensity exercise might persist at 24h follow-up, while the high intensity exercise might decrease the pain threshold in the next day. (3) The CPM responses at 24h following high intensity running might attenuated compared with the low and moderate intensity running.

## Methods

This study has been approved (2023023H) by Sports Science Experimental Ethics Committee of Beijing Sport University.

### Study design

A total of sixty-nine healthy participants were included in this study, and they were invited to perform exercise interventions with different intensities. All participants were provided with consent forms before undergoing the study. The demographic data and baseline measurements (e.g., resting-heart rate (HRrest), pressure pain threshold/tolerance, and CPM responses) were collected. The maximum-heart rate (HRmax) was estimated using the formula^30^: HRmax=202.5-0.53*age, and the reserved heart rate (HRR) was calculated as HRR=HRmax-HRrest. The real-time HR were collected and recorded via the HR belt worn by the participants during running. To avoid potential long-lasting analgesic effects of the CPM test, all exercise interventions performed one week after the baseline measurements.

All individuals who conformed to this study were randomly assigned to 3 experimental groups (A, B, and C) with different exercise intensity in a 1:1:1 ratio. And the randomized sequences were generated by the Excel software. All of the participants were labeled from number 01 to 66, and allocated following the A-B-C circulation order. The screeners of the participants were AN and XZH.

The participants would execute the low-intensity treadmill running with 40% HRR in group A, moderate-intensity (55% HRR) in group B, or high-intensity (70% HRR) in group C. And the speed of the running would be determined in coherence with the target heart rate (THR) during the baseline measurements. All participants would perform a single exercise session with predetermined intensity for 30min. (Figure 1)

**Figure 1.**
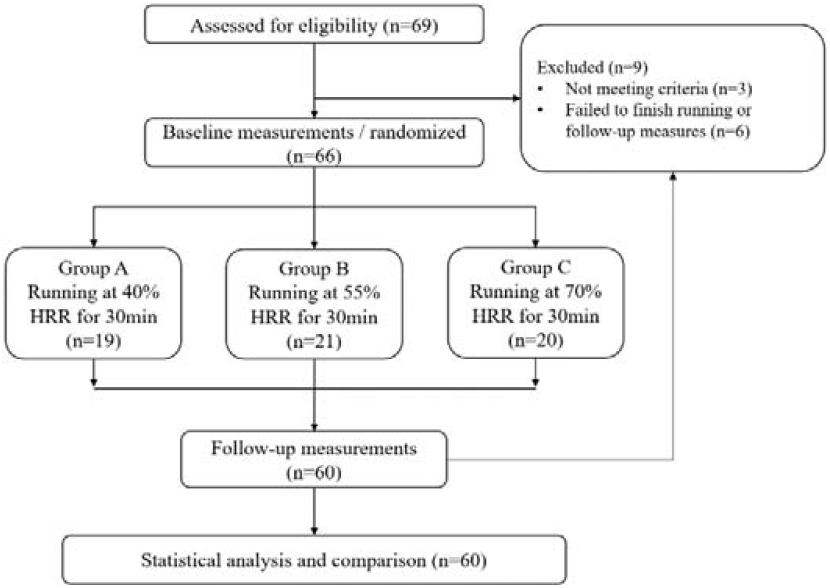
Flowchart of the experiment

### Participants

Based on the previous studies^31,32^, the aerobic exercise-induced effect size on PPT changes ranged from 0.20 to 0.38, and PPTol was 0.20. Our study utilized G-Power Software with the effect size = 0.38; alpha level = 0.05; and power = 0.80. Thus, a minimum total sample size of 66 participants across the three groups was determined.

A total of sixty-nine healthy female students (aged 18 to 30 years) from Beijing Sport University were included in this study, and 66 of them were finally enrolled. The exclusion criteria were: (1) had pain or pain-related syndrome within 3 months; (2) had injury history of lower extremities within 1 year; (3) had potential or confirmed heart disease, or recovered from a heart disease less than 1 year, (4) failed to maintain or tolerant the exercise intensity during the treadmill running; (5) showed serious exertion or fatigue in 24h after exercise sessions; (6) showed intolerable pain during the pain perception test or the CPM test.

### Procedures

All participants performed a single treadmill running session with different intensities based on their THR. The THR was 40% HRR in group A, 55% HRR in group B, and 70% HRR in group C. Participants worn an HR belt to monitor and record real-time HR during the test and running session. (Figure 2)

**Figure 2.**
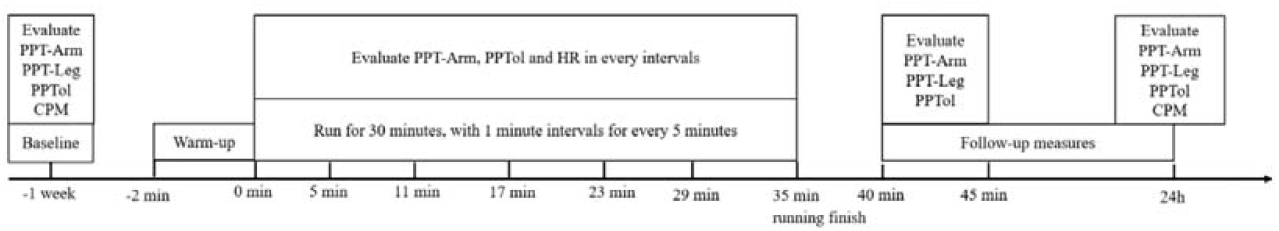
Flowchart of the procedures

The baseline PPT and PPTol were measured 10 min before the exercise session, and the CPM responses were tested in baseline measurement to mitigate the possibility of long-lasting analgesic effects. The exercise session consisted of 30min running with 5 interval periods (at the speed of 4 km/h for 1 min) in every 5min (35min in total). Throughout each interval period, heart rate (HR), PPT-arm, and PPTol were measured. Post-exercise PPT and PPTol were conducted at 5min and 10min after the exercise. Additionally, PPT, PPTol and CPM were measured 24h after the exercise session.

### Outcome measures

The outcome measures were assessed at multiple time points: before, during, and after the running session, where the PPT-arm and PPTol were recorded in every interval during running, and the PPT-leg were only tested after running session. And the CPM responses were evaluated by cold pressure methods during baseline session and 24h after the running session.

#### Pressure pain threshold (PPT)

The PPT was evaluated by a quantitative sensory testing protocol^33^ with a handheld pressure algometer (Baseline Dolorimeter, Fabrication Enterprises, USA) equipped with a 1 cm^2^ metal probe. The pressure was applied at a rate of 0.5kg/s over two locations: the extensor carpus radialis (PPT-arm) and the peroneus longus (PPT-leg) on the right side. Participants were instructed to indicate their perceived pain intensity using a visual analog score (VAS) ranging from 0 to 100. When participants reported a pain intensity of 30 out of 100 (Pain30) during the pressure application, the pressure thresholds were recorded as the PPT values.

#### Pressure pain tolerance (PPTol)

The PPTol was assessed using a quantitative sensory testing protocol^34^ via handheld pressure algometer (Baseline Dolorimeter, Fabrication Enterprises, USA) with a 1 cm^2^ metal probe. The pressure was applied at a rate of 0.5kg/s over the extensor carpus radialis on the left side. Participants were instructed to indicate their perceived pain intensity using a visual analog score (VAS) ranging from 0 to 100. When participants reported a pain intensity of 70 out of 100 (Pain70) during the pressure application, the pressure threshold was recorded as the PPTol value.

#### Conditioned pain modulation (CPM)

The CPM response was measured using a quantitative sensory testing protocol, specifically employing the cold pressor procedure^35^, In this procedure, pressure was applied as the test stimulation, while cold water immersion served as the conditioned stimulation. Participants first received pressure stimulation at the ipsilateral extensor carpus radialis and reported the PPT when the pain intensity reached Pain30 as a test stimulus. Subsequently, participants were instructed to immerse the contralateral hand into cold water at 8□ for 1 min. The PPT at Pain30 was reassessed when participants withdrew their hand from the immersion. The difference between the two PPTs was recorded as the response of CPM.

### Statistical analysis

The normality of all data was assessed using the Shapiro-Wilk test. The difference in baseline data (height, weight, HRrest, CPM, PPT and PPTol) between groups were analyzed using a one-way analysis of variance (ANOVA) test. To determine the differences within the three groups over time (running times and the acute follow-up times), a two-way (running time and intensity) repeated measures ANOVA was applied to examine PPT and PPTol, except for the PPT-leg at 35min, which were tested via the one-way analysis of covariance (ANCOVA) with the baseline measurements set as covariates. For the changes in PPT and PPTol at 24h post-running, the one-way ANCOVA method were also applied. The changes of CPM responses were evaluated by the one-way ANCOVA, while the baseline measurements were set as covariates. Post-hoc multiple comparisons were conducted using the Bonferroni method. All statistical analyses were performed using SPSS Version 21.0, and a significance level of p<0.05 was applied to all tests.

## Results

### Baseline and running characteristics

Three participants were excluded from this study for the shoulder pain syndrome occurred 1 month before the experiments. Of the sixty-six participants enrolled in this study, 19 of them in group A completed running for 30min with low intensity, 21 of them in group B completed running for 30min with moderate intensity, 20 of group C completed running for 30min with high intensity. And 6 participants did not complete running sessions or follow-up measurements and dropped out of this study. All of baseline characteristics did not present significant differences between the groups (p > 0.05, Table 1). There were significant differences in HR for every interval among groups during the running periods (Table 1).

**Table 1.** Baseline and running measurement (M±SD) ^1^

| Measurements | A (n=19) | B(n=21) | C(n=20) | P |
| --- | --- | --- | --- | --- |
| Age (y) | 21.63±2.01 | 22.52±2.06 | 21.10±2.20 | 0.097 |
| Height (cm) | 166.95±7.31 | 166.48±7.27 | 165.55±4.93 | 0.797 |
| Weight (kg) | 59.05±9.67 | 56.05±9.23 | 57.55±8.02 | 0.576 |
| PPT-Arm (kg/cm<sup>2</sup>) | 2.31±0.16 | 2.33±0.13 | 2.32±0.17 | 0.949 |
| PPT-Leg (kg/cm<sup>2</sup>) | 4.40±0.53 | 4.51±0.48 | 4.46±0.48 | 0.799 |
| PPTol (kg/cm<sup>2</sup>) | 4.57±0.42 | 4.68±0.49 | 4.61±0.40 | 0.740 |
| CPM (kg/cm<sup>2</sup>) | 0.70±0.15 | 0.70±0.16 | 0.71±0.17 | 0.998 |
| HRrest (beats/min) | 81.11±0.89 | 82.71±6.86 | 81.95±7.79 | 0.814 |
| THR (beats/min) | 125.26±5.32 | 142.03±3.07 | 158.51±2.40 | <0.001 |
| HR of 5min (beats/min) | 125.85±5.75 | 143.95±5.96 | 161.00±4.67 | <0.001 |
| HR of 11min (beats/min) | 126.36±6.01 | 143.38±6.67 | 163.35±3.92 | <0.001 |
| HR of 17min (beats/min) | 126.00±6.45 | 142.62±5.69 | 164.00±4.34 | <0.001 |
| HR of 23min (beats/min) | 125.32±6.80 | 143.95±5.98 | 165.01±4.52 | <0.001 |
| HR of 29min (beats/min) | 126.11±6.43 | 143.71±4.66 | 165.95±3.79 | <0.001 |
| HR of 35min (beats/min) | 126.47±5.86 | 144.14±5.34 | 166.15±4.15 | <0.001 |

<sup>1</sup>: 1-way ANOVA, significant difference was set by $p \leq 0.05$ .

### Changes in PPT of arm following running

The two-way repeated-measures ANOVA revealed significant main effects (F=264.74, p<0.001) for the running time on the PPT of the arm, which indicated that running for 30min significantly increased global PPT. The interaction effect (F=13.55, p<0.001) between running intensity and time on the PPT of the arms was also significant. Post-hoc comparisons showed that the changes of PPT in moderate intensity group were significantly higher than both the low (p=0.003) and high (p<0.001) intensity groups. Furthermore, the low intensity running also resulted in a higher PPT (p<0.001) compared to the high intensity running.

The two-way repeated-measures ANOVA showed significant main effects (F=385.83, p<0.001) for the running time on the PPT of the arm at immediate post-running. The interaction effect (F=25.35, p<0.001) of the running intensity and time point on the PPT of the arm was also significant. Post-hoc comparisons showed that the changes in PPT for high intensity group were significantly lower (p<0.001) compared to other two groups. However, there were no significant differences between the low and moderate intensity groups (p=0.09) in terms of their PPT changes.

The one-way ANCOVA revealed significant between-groups differences (F=16.52, p<0.001) at 24-hour follow-up when considering the running intensities. Post-hoc comparisons showed that the changes in PPT for high intensity group were significantly lower (p<0.001) compared to other two groups, but no significant differences between low and moderate groups (p=0.80). (Table 2, Table 3 and Figure 3)

**Figure 3.**
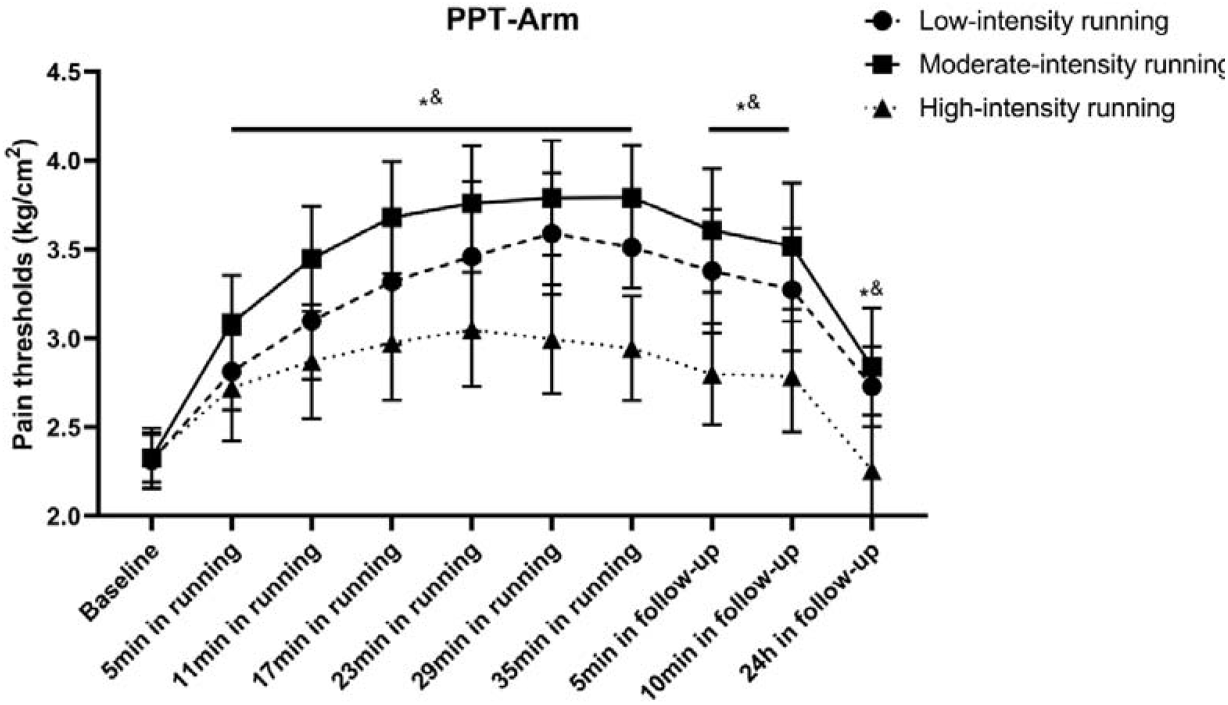
Changes in PPT of arms following running *: PPT in low intensity group significantly higher than high intensity group ^&^: PPT in moderate intensity group significantly higher than high intensity group

### Changes in PPT of leg following running

The two-way ANCOVA revealed significant between-groups differences (F=19.25, p<0.001) on the PPT of the legs at immediate post-running. Post-hoc comparisons showed that the changes in PPT for high intensity group were significantly lower than both the low (p=0.002) and moderate (p<0.001) intensity groups. However, there were no significant differences between low and moderate intensity groups.

The two-way repeated-measures ANOVA showed significant main effects (F=181.03, p<0.001) for the measurement time on the PPT of the legs at 5-10min post-running. The interaction effect (F=18.65, p<0.001) of the running intensity and time on the PPT of the legs was also significant. Post-hoc comparisons showed that the changes in PPT for high intensity group were significantly lower than both the low (p=0.007) and moderate (p<0.001) intensity groups.

The one-way ANCOVA revealed significant between-groups differences (F=38.11, p<0.001) at 24-hour follow-up when considering the running intensities. Post-hoc comparisons showed that the changes in PPT for high intensity group were significantly lower (p<0.001) compared to other two groups, but no significant differences between low and moderate groups (p>0.999). (Table 4, Table 5 and Figure 4)

**Figure 4.**
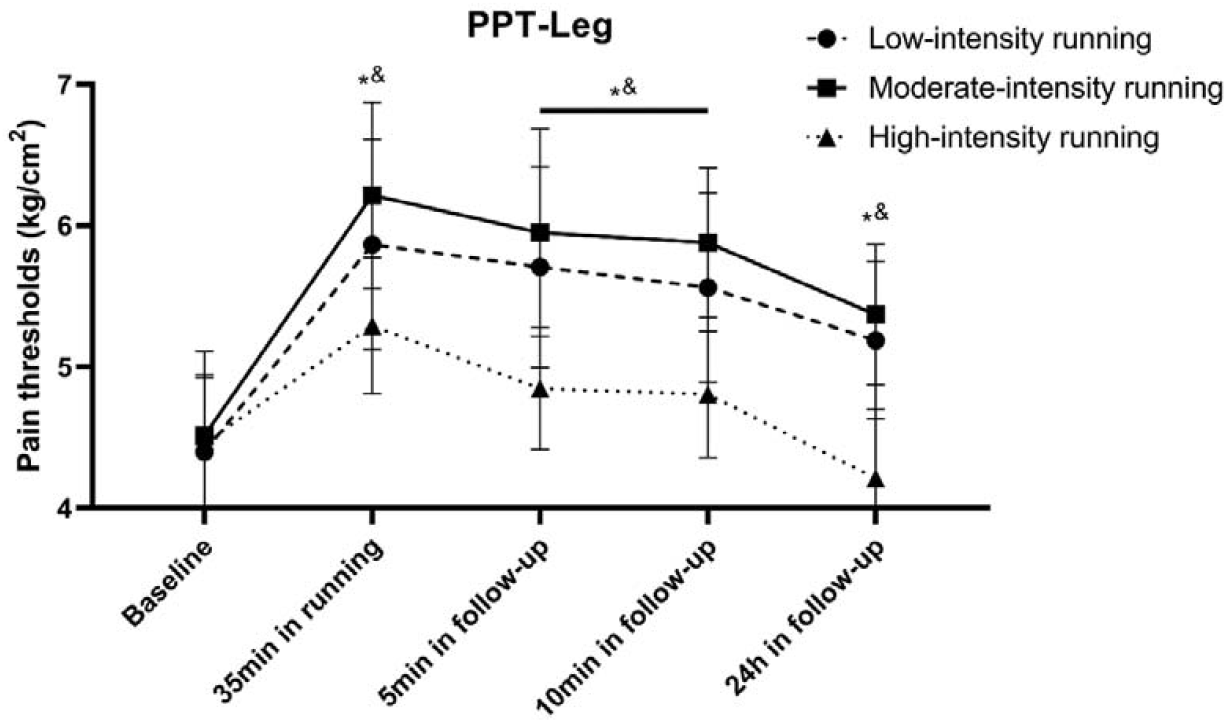
Changes of PPT in legs following running *: PPT in low intensity group significantly higher than high intensity group ^&^: PPT in moderate intensity group significantly higher than high intensity group

### Changes in PPTol following running

The two-way repeated-measures ANOVA revealed significant main effects (F=122.93, p<0.001) for the running time on the PPTol, which indicated that the running for 30min significantly increased PPTol. The interaction effect (F=3.90, p<0.001) between running intensity and time on the PPTol was also significant. Post-hoc comparisons showed that the changes in PPTol for moderate intensity group were significantly higher (p=0.008) than high intensity groups. However, there were no significant differences between low and high intensity groups (p=0.129), or low and moderate intensity groups (p=0.256).

The two-way repeated-measures ANOVA showed significant main effects (F=150.96, p<0.001) for the measurement time on the PPTol at 5-10min post-running. The interaction effect (F=4.18, p=0.02) between running intensity and time on the PPTol was also significant. Post-hoc comparisons showed that the changes in PPTol for moderate intensity group were significantly higher (p=0.006) than high intensity groups. However, there were no significant differences between low and high intensity groups (p=0.140), or low and moderate intensity groups (p=0.199).

The one-way ANCOVA revealed significant between-groups differences (F=13.58, p<0.001) at 24-hour follow-up when considering the running intensities. Post-hoc comparisons showed that the changes in PPTol for high intensity group were significantly lower (p<0.001) compared to other two groups, but no significant differences between low and moderate intensity groups (p>0.999). (Table 6, Table 7 and Figure 5)

**Figure 5.**
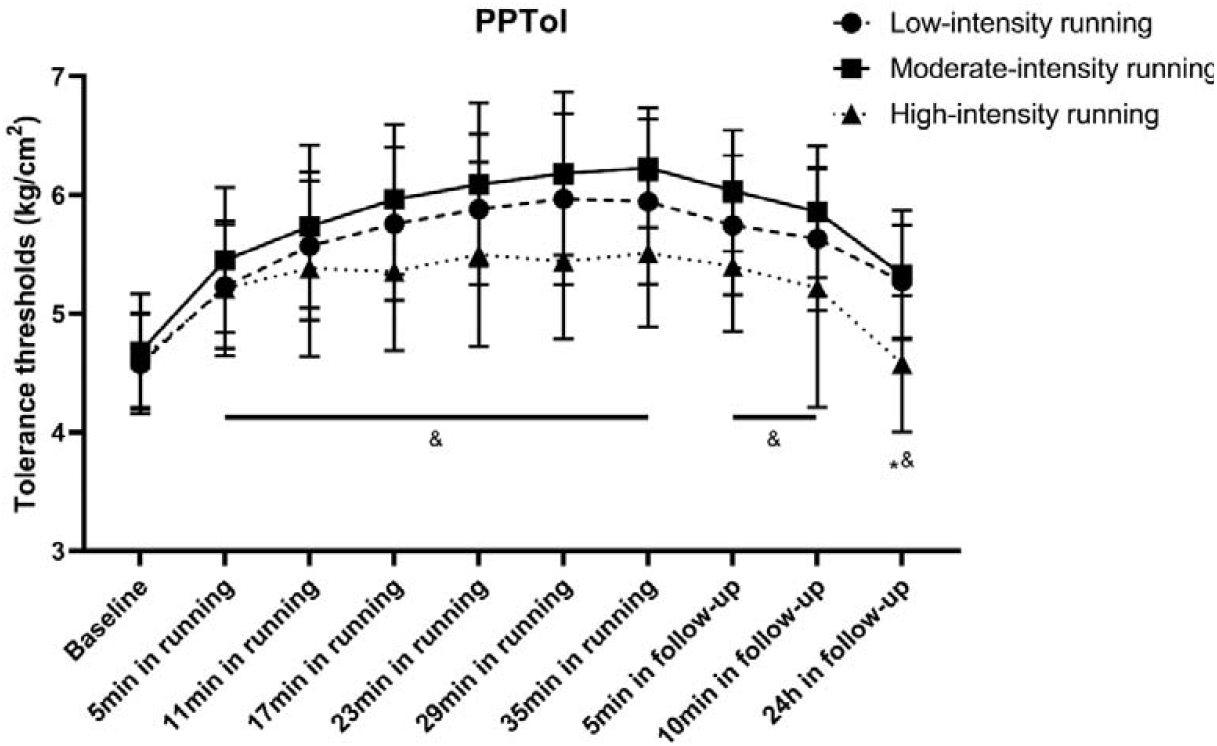
Changes in PPTol following running *: PPTol in low intensity group significantly higher than high intensity group ^&^: PPTol in moderate intensity group significantly higher than high intensity group

### Changes in CPM following running

The one-way ANCOVA revealed significant between-groups differences (F=27.17, p<0.001) at 24-hour follow-up when considering the running intensities. Post-hoc comparisons showed that the changes in CPM for high intensity group were significantly lower (p<0.001) compared to other two groups, but no significant differences between low and moderate groups (p>0.999). (Table 8, Table 9 and Figure 6)

**Figure 6.**
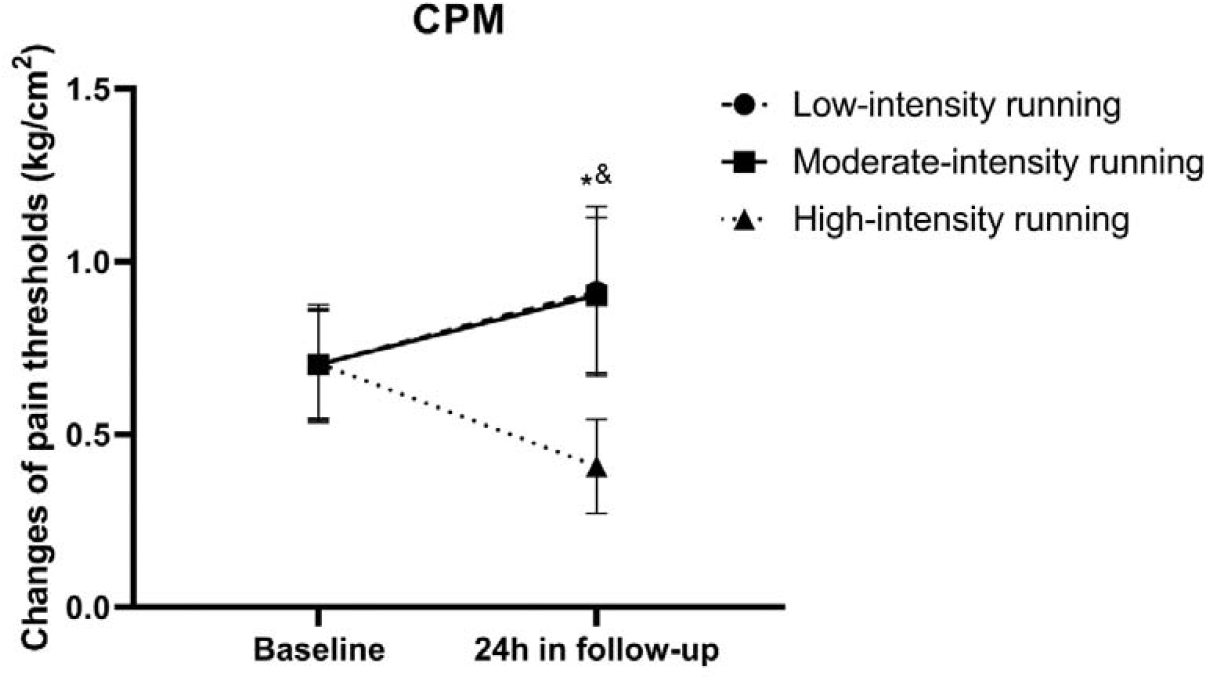
Changes in CPM following running *: CPM in low intensity group significantly higher than high intensity group ^&^: CPM in moderate intensity group significantly higher than high intensity group

## Discussion

This study aimed to investigate the modulation of EIH effects following running exercises with different intensities in healthy individuals. Our results showed that the changes of PPT and PPTol had the temporal summation effects, which both increased along with the running time. The PPT and CPM responses of the moderate and low intensity running were significantly higher than the exercise with high intensity during running sessions and follow-ups. Furthermore, the improvements of PPTol following moderate intensity running were also significantly greater than high intensity running. These results indicated that the modulation of EIH effects may involve distinct central mechanisms that are influenced by the context of the exercise.

Previous studies have examined the changes in PPT among healthy individuals following aerobic exercise with different intensities. For instance, Hoffman et al^36^ compared the PPT changes during the running and found that running with 75% of maximal oxygen consumption (VO_2_max) elicited greater EIH effects than the running of 50% VO_2_max. And in contrast, Kruger et al^27^ observed that the maximal endurance cycling induced mechanical allodynia. A cross-over study by Hviid et al^32^ showed that fast walking over 60 mins can increase the PPT significantly. These results indicated that the EIH effect may not increase along with the exercise intensities. However, most of them only evaluated acute EIH effects, and did not investigate the secondary changes at 24h following running, which were insufficient to reveal the role of endogenous pain modulation in the EIH effects.

For the reason of different EIH effects following exercises with various intensities, it may be explained by the interaction between the muscle contraction and the C fibers inputs during the treadmill running. Muscle contraction can activate the C fibers during the exercise^17^. Thus, the high-intensity exercises may trigger the descending inhibition, and upregulate the opioids^37,38^ in PAG. However, it can also activate noxious C fibers, potentially inducing descending facilitation and ultimately decreasing EIH and CPM responses. While the moderate and low intensity exercises with sufficient time may stimulate non-noxious C fibers, triggering the activation of cannabinoid^39^ and 5-HT^5,40^ receptors in the PAG and RVM. This activation can enhance the CPM responses and analgesic effects. (Figure 7)

**Figure 7.**
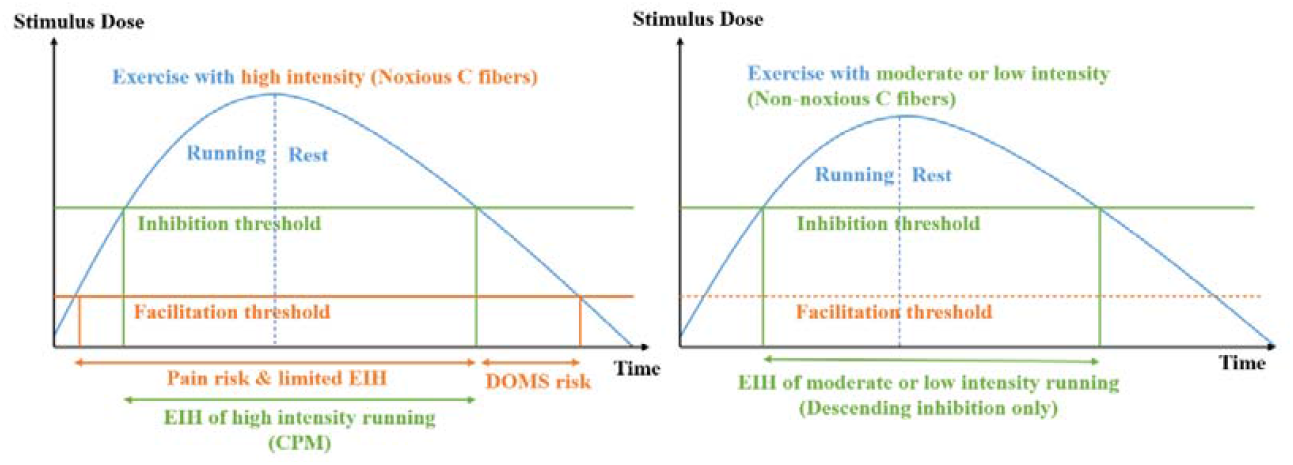
Potential mechanisms of EIH induced via running with different intensities EIH: exercise-induced hypoalgesia; DOMS: delayed onset muscle soreness; CPM: conditioned pain modulation.

The role of CPM responses in EIH effects has also been investigated by Lemley et al^41^. They compared EIH responses in healthy individuals based on CPM levels, and found that adults with greater CPM were more likely to experience greater EIH, which were also consistent with the findings in our previous study^42^. Thus, the descending inhibition might contribute to the PPT and PPTol changes following the aerobic exercises in our study, which might be modulated by the intensities of exercises in healthy individuals.

Additionally, it has been observed that low or moderate intensity exercises can improve pain tolerance in individuals with conditions. Vaegter et al^43,44^ investigated the EIH responses during aerobic and resistance exercises and found that the increase in pain tolerance following low-intensity exercises occurred in both healthy and pain individuals. Hviid et al^32^ also found that the changes in PPTol can be elicited by walking exercise in relatively low intensity.

The changes of pain tolerance observed during exercise may also be attributed to the improvements in cognitive process^6^ related to pain perception in brain areas of “Pain Matrix”^45^ These areas include the prefrontal cortex^46^, ventral tegmental area (VTA)^47^, amygdala^8^, nucleus accumbens (NAc), hippocampus and insular^48^. Some rodent studies on neuropathic pain^49,50^ reported that the voluntary exercise activated the dopamine neurons in the VTA and the gamma-aminobutyric acid neurons in the NAc shell, which are associated with pain tolerance and emotional aspects such as anxiety, depression and fear related to pain perception^51,52^. Furthermore, Baiamonte et al^9^ found that a negative relationship between changes in PPTol during resistance exercises and HR and rating of perceived exertion levels, indicating that exercise fatigue may attenuate the EIH effects and reduce tolerance to pain perception.

There were several limits in this study. First, the indicators of the pain tests were limited, for instance, adding the heat pain detection and tolerance thresholds might give a more complete description of the changes in pain perception. Second, the results of individuals’ pain tolerance thresholds might can be influenced by previous pain experiences and subjective emotional perceptions, leading to a diverse range of responses. Lastly, all of the participants in this study were female. Considering the potential gender differences in EIH and CPM responses following running exercise, future studies should take into account the inclusion of both male and female participants to provide a comprehensive understanding of these effects

## Conclusion

The finding of our study suggested that the running with moderate and low intensities induced more pronounced and longer lasting EIH effects in healthy individuals than high intensity running, which might be modulated by the endogenous descending pain modulations considering the context of exercises. The changes of PPT, PPTol and CPM indicated that the high-intensity running may activate the descending facilitation and inhibition effects, hereby limiting the efficacy of endogenous analgesia and attenuating CPM responses. While the moderate and low intensity exercise with sufficient time can also activate the endogenous descending inhibition pathway, leading to a reduction in pain perception. Thus, the pain perception changes following exercises may reveal the potential mechanisms of EIH, which can be considered in the practice of chronic pain managements and future researches.

## Acknowledgments

We would like to thank Xin Wang, Dong-Mei Yu, Na Zhang, Shuo-Yan Wang and all the researchers who provided help and advice in our experiments.

## Supplemental files

**Table 2.** Changes in PPT-Arm following running exercise (M±SD)

| Measurements | A (n=19) | B(n=21) | C(n=20) | P |
| --- | --- | --- | --- | --- |
| Baseline | 2.31±0.16 | 2.33±0.13 | 2.32±0.17 |  |
| 5min in running<sup>1</sup> | 2.81±0.21 | 3.09±0.27 | 2.72±0.30 |  |
| 11min in running<sup>1</sup> | 3.10±0.33 | 3.45±0.30 | 2.87±0.32 |  |
| 17min in running<sup>1</sup> | 3.32±0.35 | 3.68±0.31 | 2.98±0.32 | <0.001<sup>1</sup> |
| 23min in running<sup>1</sup> | 3.46±0.42 | 3.78±0.32 | 3.05±0.32 |  |
| 29min in running<sup>1</sup> | 3.60±0.34 | 3.80±0.32 | 3.00±0.31 |  |
| 35min in running<sup>1</sup> | 3.51±0.23 | 3.79±0.29 | 2.95±0.29 |  |
| 5min in follow-up<sup>2</sup> | 3.38±0.35 | 3.61±0.35 | 2.80±0.28 | <0.001<sup>2</sup> |
| 10min in follow-up<sup>2</sup> | 3.27±0.34 | 3.52±0.36 | 2.79±0.31 |  |
| 24h in follow-up<sup>3</sup> | 2.73±0.23 | 2.83±0.33 | 2.26±0.31 | <0.001<sup>3</sup> |

<sup>1</sup>: 2-way Repeated measures ANOVA, significant difference was set by $p \leq 0.05$ .<sup>2</sup>: 2-way Repeated measures ANOVA, significant difference was set by $p \leq 0.05$ .<sup>3</sup>: 1-way ANCOVA, significant difference was set by $p \leq 0.05$ .

**Table 3.** Between-group comparison results of PPT-Arm following running exercise

| Measurements | Groups |  | Mean Difference | Std. Error | P | 95% CI |  |
| --- | --- | --- | --- | --- | --- | --- | --- |
|  |  |  |  |  |  | Lower Bound | Upper Bound |
| 0-35min in running<sup>1</sup> | A | B | -0.254 | 0.073 | 0.003 | -0.435 | -0.073 |
|  |  | C | 0.317 | 0.074 | <0.001 | 0.134 | 0.500 |
|  | B | C | 0.571 | 0.072 | <0.001 | 0.393 | 0.750 |
| 5-10min in follow-up<sup>2</sup> | A | B | -0.162 | 0.073 | 0.091 | -0.342 | 0.018 |
|  |  | C | 0.351 | 0.074 | <0.001 | 0.169 | 0.533 |
|  | B | C | 0.513 | 0.072 | <0.001 | 0.336 | 0.691 |
| 24h follow-up<sup>3</sup> | A | B | -0.103 | 0.092 | 0.800 | -0.329 | 0.123 |
|  |  | C | 0.478 | 0.093 | <0.001 | 0.249 | 0.707 |
|  | B | C | 0.581 | 0.090 | <0.001 | 0.358 | 0.804 |

<sup>1</sup>: 2-way Repeated measures ANOVA, adjusted by Bonferroni. significant difference was set by $p \leq 0.05$ .<sup>2</sup>: 2-way Repeated measures ANOVA, adjusted by Bonferroni. significant difference was set by $p \leq 0.05$ .<sup>3</sup>: 1-way ANCOVA, adjusted by Bonferroni. significant difference was set by $p \leq 0.05$ .

**Table 4.**
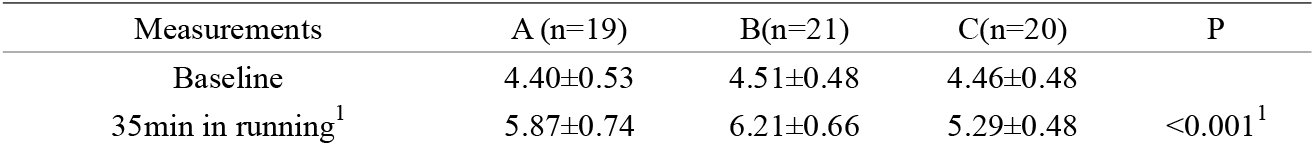
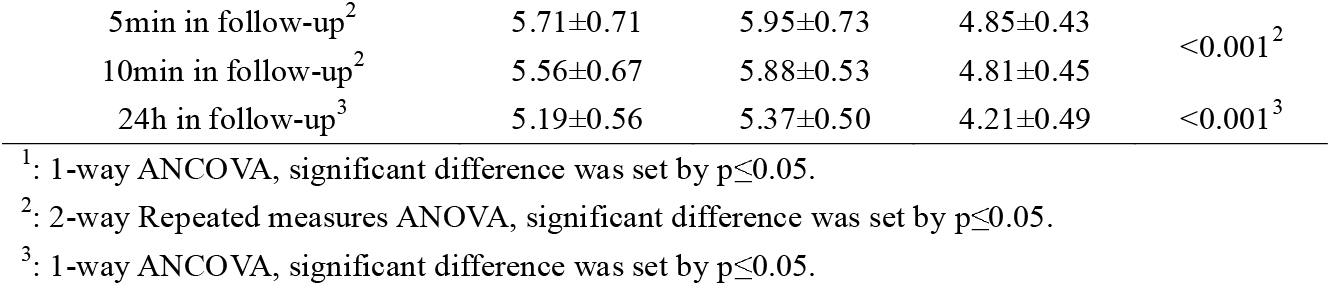
Changes in PPT-Leg following running exercise (M±SD)

**Table 5.** Between-group comparison results of PPT-Leg following running exercise

| Measurements | Groups |  | Mean Difference | Std. Error | P | 95% CI |  |
| --- | --- | --- | --- | --- | --- | --- | --- |
|  |  |  |  |  |  | Lower Bound | Upper Bound |
| 0-35min in running<sup>1</sup> | A | B | -0.272 | 0.169 | 0.336 | -0.688 | 0.144 |
|  |  | C | 0.614 | 0.170 | 0.002 | 0.195 | 1.034 |
|  | B | C | 0.886 | 0.166 | <0.001 | 0.477 | 1.296 |
| 5-10min in follow-up<sup>2</sup> | A | B | -0.226 | 0.162 | 0.507 | -0.626 | 0.174 |
|  |  | C | 0.519 | 0.164 | 0.007 | 0.115 | 0.924 |
|  | B | C | 0.745 | 0.160 | <0.001 | 0.351 | 1.140 |
| 24h follow-up<sup>3</sup> | A | B | -0.119 | 0.135 | >0.999 | -0.453 | 0.214 |
|  |  | C | 1.009 | 0.136 | <0.001 | 0.672 | 1.345 |
|  | B | C | 1.128 | 0.133 | <0.001 | 0.800 | 1.456 |

<sup>1</sup>: 1-way ANCOVA, adjusted by Bonferroni. significant difference was set by $p \leq 0.05$ .
<sup>2</sup>: 2-way Repeated measures ANOVA, adjusted by Bonferroni. significant difference was set by $p \leq 0.05$ .
<sup>3</sup>: 1-way ANCOVA, adjusted by Bonferroni. significant difference was set by $p \leq 0.05$ .

**Table 6.** Changes in PPTol following running exercise (M±SD)

| Measurements | A (n=19) | B (n=21) | C (n=20) | P |
| --- | --- | --- | --- | --- |
| Baseline | 4.57±0.42 | 4.68±0.49 | 4.61±0.40 |  |
| 5min in running<sup>1</sup> | 5.23±0.52 | 5.45±0.61 | 5.21±0.57 |  |
| 11min in running<sup>1</sup> | 5.57±0.62 | 5.73±0.69 | 5.38±0.74 |  |
| 17min in running<sup>1</sup> | 5.76±0.64 | 5.96±0.63 | 5.36±0.67 | <0.001<sup>1</sup> |
| 23min in running<sup>1</sup> | 5.88±0.63 | 6.09±0.69 | 5.50±0.78 |  |
| 29min in running<sup>1</sup> | 5.97±0.72 | 6.18±0.69 | 5.44±0.66 |  |
| 35min in running<sup>1</sup> | 5.94±0.69 | 6.23±0.50 | 5.51±0.62 |  |
| 5min in follow-up<sup>2</sup> | 5.74±0.59 | 6.04±0.51 | 5.40±0.55 | <0.001<sup>2</sup> |
| 10min in follow-up<sup>2</sup> | 5.63±0.60 | 5.86±0.56 | 5.22±1.00 |  |
| 24h in follow-up<sup>3</sup> | 5.27±0.47 | 5.33±0.54 | 4.58±0.57 | <0.001<sup>3</sup> |

<sup>1</sup>: 2-way Repeated measures ANOVA, significant difference was set by $p \leq 0.05$ .
<sup>2</sup>: 2-way Repeated measures ANOVA, significant difference was set by $p \leq 0.05$ .
<sup>3</sup>: 1-way ANCOVA, significant difference was set by $p \leq 0.05$ .

**Table 7.** Between-group comparison results of PPTol following running exercise

| Measurements | Groups | Mean | Std. Error | P | 95% CI |
| --- | --- | --- | --- | --- | --- |

|  |  |  | Difference |  |  | Lower Bound | Upper Bound |
| --- | --- | --- | --- | --- | --- | --- | --- |
| 0-35min in running<sup>1</sup> | A | B | -0.201 | 0.175 | 0.256 | -0.552 | 0.150 |
|  |  | C | 0.273 | 0.177 | 0.129 | -0.082 | 0.628 |
|  | B | C | 0.474 | 0.173 | 0.008 | 0.128 | 0.821 |
| 5-10min in follow-up<sup>2</sup> | A | B | -0.208 | 0.160 | 0.199 | -0.529 | 0.113 |
|  |  | C | 0.243 | 0.162 | 0.140 | -0.082 | 0.567 |
|  | B | C | 0.451 | 0.158 | 0.006 | 0.134 | 0.767 |
| 24h follow-up<sup>3</sup> | A | B | -0.002 | 0.156 | >0.999 | -0.387 | 0.382 |
|  |  | C | 0.711 | 0.157 | <0.001 | 0.323 | 1.098 |
|  | B | C | 0.713 | 0.153 | <0.001 | 0.335 | 1.091 |

<sup>1</sup>: 2-way Repeated measures ANOVA, adjusted by Bonferroni. significant difference was set by $p \leq 0.05$ .
<sup>2</sup>: 2-way Repeated measures ANOVA, adjusted by Bonferroni. significant difference was set by $p \leq 0.05$ .
<sup>3</sup>: 1-way ANCOVA, adjusted by Bonferroni. significant difference was set by $p \leq 0.05$ .

**Table 8.** Changes in CPM following running exercise (M±SD)

| Measurements | A (n=19) | B(n=21) | C(n=20) | P |
| --- | --- | --- | --- | --- |
| Baseline | 0.70±0.15 | 0.70±0.16 | 0.71±0.17 |  |
| 24h in follow-up<sup>1</sup> | 0.91±0.25 | 0.90±0.22 | 0.41±0.14 | <0.001<sup>1</sup> |

<sup>1</sup>: 1-way ANCOVA, significant difference was set by $p \leq 0.05$ .

**Table 9.** Between-group comparison results of CPM following running exercise^1^

| Groups |  | Mean Difference | Std. Error | Sig. | 95% CI |  |
| --- | --- | --- | --- | --- | --- | --- |
|  |  |  |  |  | Lower Bound | Upper Bound |
| A | B | 0.011 | 0.065 | >0.999 | -0.150 | 0.171 |
|  | C | 0.506 | 0.066 | <0.001 | 0.344 | 0.668 |
| B | C | 0.496 | 0.064 | <0.001 | 0.337 | 0.654 |

<sup>1</sup>: 1-way ANCOVA, adjusted by Bonferroni. significant difference was set by $p \leq 0.05$ .

