## supplemental file 1 for "Modulation of Pain Perceptions Following Treadmill Running with Different Intensities and The Potential Mechanisms of Exercise-induced Hypoalgesia"

### 北京体育大学运动科学实验伦理审批表

### Ethics Approval Form for Sports Science Experiments of Beijing Sport University

编号（*No.*）:2023023H

| | | | | | |
|---|---|---|---|---|---|
| 项目名称 Study Title | 有氧运动负荷对健康人群的疼痛感受影响和内源性痛觉调控相关性的研究 | | | | |
| 研究期限 Research Period | 2023 年 5 月 30 日至 2024 年 5 月 30 日 | | | | |
| 项目负责人 Project Manager | 徐子涵 | 性别 Gender | 男 | 职称 Academic Level | 博士研究生 |
| 研究方向 Research Direction | 运动康复 | 联系方式 Phone Number | 17888837585 | 邮箱 E-mail | |
| 申请节点 Application Node | ☑ 课题申报 ☑课题开展 □其他：________ | | | | |
| 审查方式 Type of Review | ☑ 快速审查 Quick Review　□会议审查 Meeting Review　□紧急会议审查 Emergency Meeting Review | | | | |
| 审查依据 Basis of Review | ☑ 伦理审查申请书 Ethical review application<br>☑ 试验方案 Research Proposal<br>☑ 知情同意书 Informed Consent Form<br>□ 研究者简历 Curriculum Vitae of Project Manager<br>□ 安全措施及应急预案 Safety Measures and Emergency Plans<br>其他资料 Other Information ________<br>（其他资料包括：试验用品安全性资料、生产企业资质证明、试验用品提供者的资质证明） | | | | |
| 项目负责人承诺 Commitment | 以上所填内容均属实，如获批准，我将严格按照提供的方案进行研究，并遵守北京体育大学运动科学实验伦理委员会的相关规定。<br>*The above contents are true. If approved, I will conduct research in strict accordance with the provided programs and comply with the relevant regulations of Sports Science Experiment Ethics Committee of Beijing Sport University.*<br>项目负责人签字（Signature）：徐子涵 | | | | |

| | | | |
|---|---|---|---|
| **审查结果**<br>**Review Result** | ☑ 同意<br>Approve | ▫不同意<br>Disapprove | ▫作必要修改后同意<br>Agreed After Modification |
| **审查结论**<br>**Evaluation** | 根据该研究的试验设计，经伦理委员会审查，受试者的健康、权利和隐私得到充分地保护，对受试者的潜在风险和伤害可控制到最小。同意开展研究。<br><br>Based on the experimental design of the study, the subject's health, rights, and privacy were adequately protected by the ethics committee review, and the potential risks and harm to the subject were minimized. Agree to conduct the study. | | |

**伦理委员会签章：**

北京体育大学运动科学实验伦理委员会
Sports Science Experiment Ethics Committee of Beijing Sport University

日期 Date: 2023年3月2日

注：本审批表双面打印，一式一份，提交至伦理委员会签章。
